# Establishing a genetic part library for tunable double-stranded RNA circuit construction and RNA interference in *Caenorhabditis elegans*

**DOI:** 10.64898/2026.03.29.715104

**Authors:** Mingyuan Xu, Sally W. Ireri, Mark Prator, C. Phoebe Lostroh, Mengyi Cao

**Author notes:** To whom correspondence may be addressed: Mengyi Cao. Mengyi Cao current address: Department of Entomology, University of Wisconsin-Madison, Madison, WI 53706.

## Abstract

RNA interference (RNAi), one of the major molecular tools in model nematode *Ceanorhabditis elegans,* relies on feeding engineered bacteria that express double-stranded RNA (dsRNA), which modulates animal host gene expression in a programmable manner. Currently no synthetic biology toolkit exits in *C. elegans* RNAi, while dsRNA circuits construction is restricted to one type of architecture. To facilitate systematic strategies for dsRNA circuit design and expression, here, we performed strain-specific screen of synthetic promoters, based on which we developed seventeen modular genetic parts compatible for rapid assembly of dsRNA expression constructs in *Escherichia coli* HT115(DE3). We validated dsRNA production in vitro and assessed RNAi efficiency in live animals by feeding. As a proof of concept, a constitutive dsRNA circuit achieved rapid and near-complete gene knockdown, whereas a *Ptac*-driven circuit enabled tunable, partial silencing while minimizing the leakiness commonly observed in standard feeding RNAi systems. Together, this work establishes a synthetic biology toolkit for programmable dsRNA delivery, enabling precise control of RNAi outcomes from partial to complete gene silencing in *C. elegans*.

## Introduction

Engineering microbes to produce and deliver double-stranded RNA (dsRNA) enables programmable RNA interference (RNAi) in multicellular eukaryotes, providing a precise and non-invasive strategy for gene silencing through natural feeding or symbiotic interactions (1–5). This approach has become a powerful tool for functional genomics and holds significant potential for applications such as species-specific pest control and crop protection (6). The mechanism of RNAi was first discovered in the nematode *Caenorhabditis elegans* (7), in which dsRNA is cleaved into small interfering RNAs (siRNAs), guiding sequence-specific cleavage and degradation of target gene. More recently, *C. elegans* has become a model for demonstrating how engineered bacterial biocircuits can program gene expression in animal hosts (8).

Two groups have created RNAi feeding libraries based on a single genetic circuit architecture in the RNaseIII-deficient feeding strain, *E. coli* HT115 (DE3), enabling dsRNA accumulation (9–10). In this strain, RNase III (*rnc)*, is interrupted by DE3 lysogeny that carries a an IPTG-inducible promoter *LacUV5* driving T7 RNA polymerase (Figure S1A) (11). In the same strain, an L4440 plasmid carries two T7 promoters with opposing directions, driving dsRNA expression of target gene in between them (Figure S1B) (9–10). Upon IPTG induction, *LacUV5* drives T7 RNA polymerase expression, enabling transcription of dsRNA under the control of T7 promoters (11). This circuit design has proven efficient for RNAi in *C. elegans* by feeding.

However, *LacUV5* is a leaky promoter, causesing knockdown of targets even in the absence of IPTG induction. In addition, a single circuit architecture may not fulfill the needs of diverse usage of the RNAi toolkits. Consequently, there is a clear need for alternative genetic parts to enable the construction of dsRNA circuits optimized for specific bacterial carrier strains and RNAi applications.

To facilitate the strategies to build diverse circuits for dsRNA expression, in this work, we established the first part library for RNAi in *C. elegans*. We built seventeen new genetic parts compatible with the CIDAR MoClo toolkit to accommodate standardized assembly that enables rapid and hierarchical construction of circuits (12, 13). This system allows for efficient combinatorial assembly of multiple DNA fragments, facilitating high-throughput design and testing of synthetic constructs. As a proof of concept, dsRNA circuits were tested in vitro and in vivo for RNA quantification and interference, showing knockdown of *C. elegans* marker gene.

## Results and Discussion

### Strain-specific promoter screen and dsRNA part library construction

The synthetic promoters from Anderson collection have primarily been characterized in *E. coli* TG1 strain (14). Because promoter strengths could be strain-specific in living bacteria, we first determined the strengths of synthetic promoters in the RNAi feeding strain *E. coli* HT115(DE3). Using the CIDAR MoClo part library (12,13), we constructed standardized circuits to test the strengths of eight Anderson promoters (Figure 1A). We quantified these promoter strengths using the expression of a Red Fluorescent Protein (RFP) reporter during bacterial growth *in vitro*. Our data show that promoter P1d causes barely detectable RFP expression (Figure 1B). Promoters P3d, P3m, P4d and P2d result in 9-16 times more RFP than the wild type, while promoters P5d, P2m, and P1m express 36-51 times more RFP than wild-type *E. coli* HT115(DE3). Overall, the relative promoter strengths we observed are largely consistent with published data for the J23100 series in *E. coli* at both 30°C and 37°C (Figure 1B and Figure S2), particularly for strong (J23101, J23102) and weak (J23103) promoters. Minor deviations in intermediate-strength promoters (e.g., J23116) likely reflect differences in strain background and experimental conditions. Based on these data, we decided to use P1m, a strong promoter for a demonstration of constitutive dsRNA circuit construction. In addition, a promoter with intermediate strength, P4d, was used in our attempt to construct the inducible but not leaky circuit for dsRNA production.

**Figure 1.**
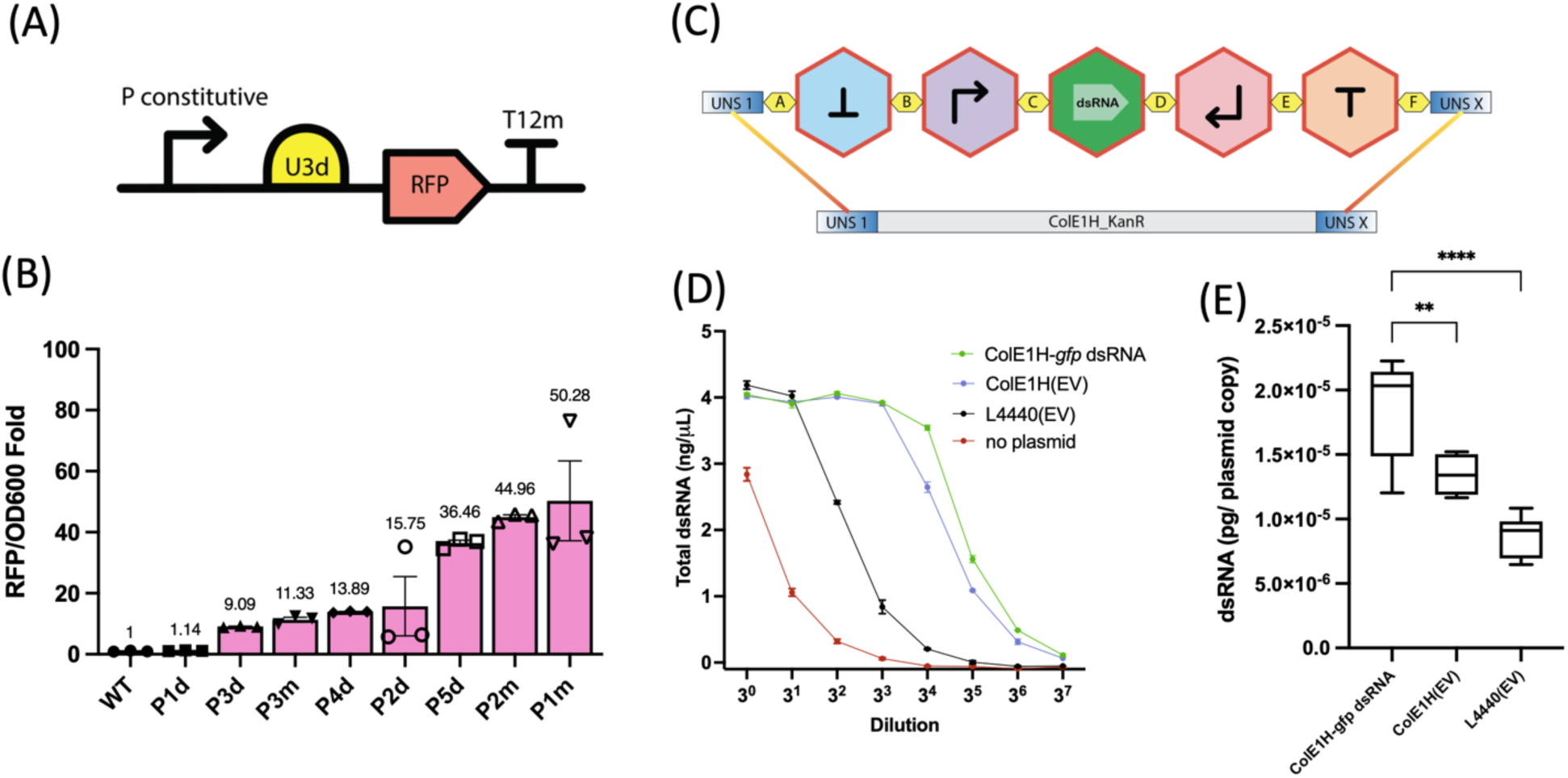
Promoter screen and dsRNA circuit construction. (A): Schematic illustration of circuit for promoter screen. (B): Synthetic promoter activities in *E. coli* HT115(DE3) assessed by RFP at 37°C. Average and standard errors of three biological replicates were shown. (C): Schematic illustration dsRNA part library and assembly strategy. Individual genetic parts were designed with CIDAR MoClo-compatible overhangs (A, B, C, D, E, and F) to enable directional assembly. UNSI and UNSX adapters to ligate transcription units into plasmid backbone. (D): dsRNA quantification assay using ELISA. *E. coli* HT115(DE3) carrying ColE1H-*gfp* dsRNA, ColE1H empty vector “EV”, L4440 empty vector, and wild-type bacterial strain without plasmid were assessed for dsRNA concentration over a serial dilution. An average and standard deviation of two biological replicates are shown for each data point. (E): Quantification of dsRNA per plasmid copy for *gfp* dsRNA expression circuit. Error bars indicate the minimum and maximum values. Statistical significance is indicated as **P < 0.01 and ****P < 0.0001.

To enable fast assembly of dsRNA circuits, we constructed seventeen new part plasmids (see Table S1) including five new types using standardized CIDAR MoClo backbone and overhang sequences for connection: reverse terminator (AB overhang), promoters (BC overhang), targeted dsRNA (CD overhang), reverse promoters (DE overhang), and terminator (EF overhang) (Figure 1C). We used v19d plasmid backbone that has a *colE1H*-like origin and is a common part used in synthetic biology. With these new part plasmids, a constitutive dsRNA expression circuit driven by P1m was created (Figure 1C, Figure 2A). As a proof of concept, expression of dsRNA driven by promoter P1m was validated by gel electrophoresis (Figure S3). Both *in vitro* transcription (IVT) of dsRNA and *E. coli* HT115(DE3) expressing P1m-driven *gfp* dsRNA produced a distinct band at the expected size of approximately 700 bp, confirming that the engineered circuit generated the correct dsRNA product. In contrast, this band was absent in the negative control, in which HT115(DE3) carried an empty vector.

**Figure 2:**
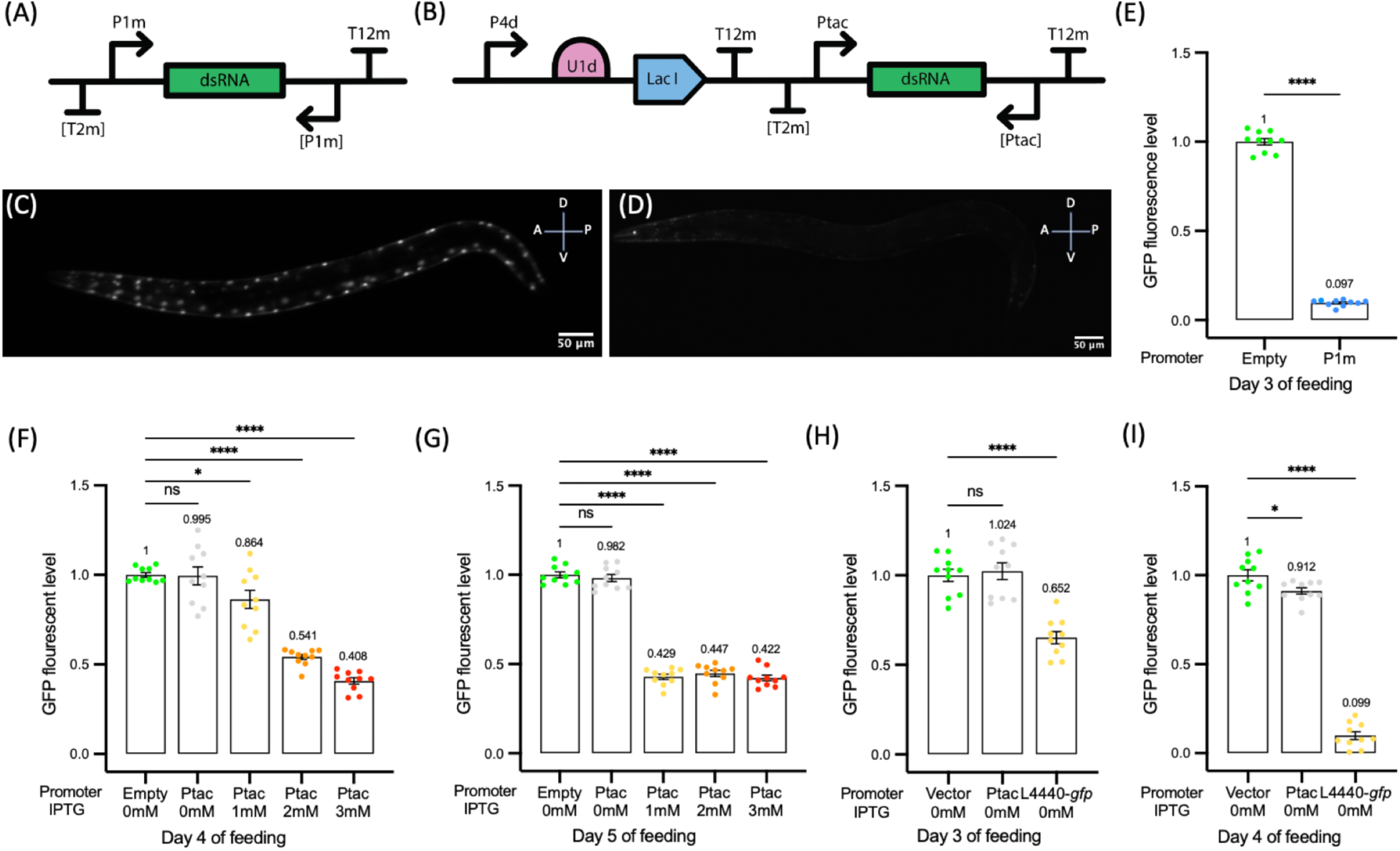
dsRNA delivery by feeding HT115(DE) in *C. elegans.* (A): Schematic illustration of a constitutive (A) and inducible (B) dsRNA circuits. (C-D): *C. elegans* (PD4251) expressing GFP feeding WT bacteria (C) or dsRNA circuit (D). “A” =anterior; “P”=posterior; “D”=dorsal’ “V”=ventral. (E): RNAi quantification in *C. elegans* feeding bacteria carrying empty plasmid (ColE1H) and constitutive dsRNA circuit. (F–G): RNAi quantification in *C. elegans* feeding on bacteria carrying control and IPTG-inducible dsRNA circuit after 4 days (F) and 5 days (G) of feeding. (H–I): RNAi quantification in *C. elegans* fed on bacteria carrying either an Ptac-based or L4440-T7-based dsRNA circuits without IPTG induction, demonstrating leakiness after 3 days (H) 4 days (I) of feeding. Each data point represents one nematode (see Methods for details of quantification). Average and standard errors are shown.

To further quantify dsRNA production, we performed a J2 antibody-based ELISA using a serial dilution series (Figure 1D). An wild-type (no plasmid) bacterial strain was used to assess the basal level of endogenous dsRNA in bacterial cells. Because the ELISA detects all dsRNA molecules longer than 15 bp, HT115(DE3) strains carrying the empty ColE1H or L4440 plasmid were also included to measure nonspecific dsRNA originating from the plasmid backbones or bacterial transcripts. As expected, the P1m-driven *gfp* dsRNA construct (ColE1H-*gfp*) produced significantly more dsRNA than all control strains (Figures 1D and 1E), yielding approximately 1.9 × 10^-5^ pg of dsRNA per plasmid copy. In comparison, the empty ColE1H and L4440 vectors produced approximately 1.3 × 10^-5^ pg and 8.6 × 10^-6^ pg of dsRNA per plasmid copy, respectively (Figure 1E). Based on our data, we estimate that approximately 28% of the total dsRNA generated by the ColE1H-*gfp* construct corresponds to the target-specific *gfp* dsRNA. Together, these results validate the functionality of the newly constructed genetic parts *in vitro* and demonstrate that the dsRNA expression circuit performs as designed.

### dsRNA delivery *in vivo* and RNAi in *C. elegans*

To ensure the sequence specificity of dsRNA in vivo, the P1m promoter-driven and constitutive dsRNA circuit (Figure 2A) was then tested for RNAi-mediated gene knockdown in *C. elegans* strain PD4251 using *gfp* as a marker (ccls4251; 15). GFP in these nematodes is found in both the body wall and vulval muscle cells and localizes to both mitochondria and nuclei (Figure 2C). As expected, feeding this nematode strain *E. coli* HT115(DE3) expressing dsRNA of *gfp* causes an RNAi-mediated knockdown of fluorescence (Figure 2D). The phenotype caused by the P1m-driven constitutive circuit expressing dsRNA was quantified (Figure 2F). After 3 days of feeding, GFP expression is ten times lower in the nematode feeding, confirming a nearly complete knockdown of GFP in the nematode cells.

Next, we constructed a Ptac-based IPTG inducible circuit for dsRNA expression and RNAi-mediated knockdown of *gfp* was tested in the same nematode strain. The circuit consists of two transcription units (TUs): TU1 includes a *lacI* driven by an intermediate strength promoter P4d (Figure 1B and 2B), while TU2 consists of a dsRNA circuit driven by the inducible promoter Ptac(LacO) (Figure 2B). Without induction, LacI is expressed moderately and constitutively to bind to Ptac(lacO), inhibiting the expression of dsRNA. Upon induction, the IPTG inducer would prevent the binding of LacI to Ptac(lacO), allowing transcription of dsRNA. As we expected, dsRNA expression and inhibition of GFP in *C. elegans* was dependent on the dose of IPTG, yielding 2.5-fold inhibition after four days of feeding with 3 mM IPTG induction (Figure 2F), confirming an inducible and partial knockdown. After five days of feeding, all three tested concentrations of IPTG resulted in about 2.5-fold RNA interference (Figure 2G).

Leakiness, or unexpected expression of dsRNA and knockdown of the host gene without IPTG induction, is a common issue with L4440 plasmid-based RNAi constructs. For this reason, we compared the IPTG-inducible Ptac-based dsRNA circuit we constructed with *colE1H* plasmid to the conventional T7-based RNAi system using L4440 plasmid. Under the same feeding conditions in the same nematode strain *E. coli* HT115(DE3) (Figure 2H and 2I). Ptac-based dsRNA circuit did not initiate RNA interference at either day 3 (Figure 2H) or day 4 (Figure 2I), indicating tight regulation of dsRNA expression without IPTG induction (not leaky). In contrast, nematodes fed with the L4440-based T7 construct showed substantial RNAi without IPTG induction, with GFP reduced to 65% at day 3 (Figure 2H) and 9.9% on day 4 (Figure 2I) relative to the vector control. These results demonstrate the characteristic leakiness of the L4440-based T7 system, whereas the Ptac-based circuit we created maintains low expression in the absence of induction.

The circuits developed here provide a generalizable framework for constructing strain-specific promoter screen and RNAi libraries in bacteria. Future work can use this platform to optimize dsRNA expression across different bacterial carrier strains, which may affect RNAi efficiency due to strain-specific RNA stability, metabolic burden, and nuclease activity (16). The modular part library, as demonstrated in this work, will accelerate extending and validating this library in a broader range of microbial carriers and eukaryotic host contexts.

### Conclusion

We performed a strain-specific promoter screen in *E. coli* HT115(DE3), based on which we constructed the first part library for *C. elegans* RNAi. Using these parts, we synthesized constitutive and inducible dsRNA that showed programmable RNA interference in host animals, overcoming the limitations of previously used RNAi libraries in *C. elegans,* and has potential to be extended to other systems.

## Methods

### Bacterial and nematode strains

*Escherichia coli* HT115(DE3) and JM109 was grown on Lysogeny Broth (LB) media supplemented with kanamycin (50 µg/mL) or carbenicillin (100 µg/mL) to maintain plasmid constructs. The *C. elegans* strain used in this study was N2 (wild-type) and PD4251 [ccIs4251 I; dpy-20(e1282) IV], obtained from the *Caenorhabditis Genetics Center* (CGC, University of Minnesota). Nematodes were propagated on the RNase-deficient cells of *E. coli* HT115(DE3) on NGM media with standard protocols for maintenance (17).

### 3G cloning and circuit construction

#### Promoter screen in *E. coli* HT115(DE3)

Transcription units containing a combination of <u>Anderson promoters</u>, RBS, and CDS (fluorescence reporter), and terminators were assembled and ligated via Golden Gate cloning (New England Biolabs) using standardized part library from CIDAR MoClo kit (12, 13), and transformed into *E. coli* HT115(DE3). To assess promoter activities in this bacterial strain, three biological replicates were used per bacterial strain (circuit), and fluorescence intensity and optical density (OD600) were acquired using BioTek Synergy H1 multimode reader (Aligent). Fluorescence measurements were normalized to cell density by calculating the ratio of fluorescence intensity to OD600 at the end time point of 24 hours growth, either at 30℃ or 37℃, providing a quantitative measure of promoter activities. Fluorescence intensities were calculated using the formula below and normalized to the WT, with the WT value set to 1.

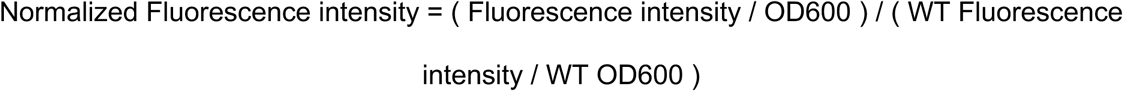

#### dsRNA circuit construction

All genetic constructs used for dsRNA circuits were designed and assembled by adapting the Golden-Gate Gibson (3G) MoClo pipeline (18). Briefly, we designed five new types of genetic parts each flanked with unique 4 bp overhang sequences (A to F): reverse-complement of transcriptional terminators (AB), promoters (BC), the RNAi targeted sequence (CD), reverse complement of promoters (DE), and transcriptional terminators (EF). A modular part library comprising 17 of these genetic elements was generated and is summarized in Table S1. Each part was cloned into an individual DVA vector and assembled into transcriptional units using Golden Gate cloning. Multiple transcriptional units were immediately assembled using compatible UNS1, UNS3, and UNS10 adaptor sequences and ligated into the ColeE1H (KanR), a linearized backbone from plasmid *V19d* by Gibson Assembly (New England Biolabs). All assembled plasmids were confirmed by whole plasmid sequencing (Plasmidsaurus) and transformed into *E. coli* HT115(DE3) for downstream dsRNA expression and RNAi experiments. All part plasmids and circuit plasmids generated in this study including plasmid map have been deposited at Addgene and are available under the plasmid IDs listed in Table S1 and Table S2.

#### dsRNA visualization assay

*E. coli* strains were prepared similarly to RNAi feeding experiments. Briefly, bacterial strains were grown in LB liquid media supplemented with kanamycin (50 µg/mL) overnight at 37℃ with aeration and then seeded onto NGM agar supplemented with kanamycin (50 µg/mL). After two days incubation at 25℃, bacteria were scraped off the NGM agar and resuspended in LB media. The collected bacterial cells were then pelleted by centrifugation and flash-frozen in liquid nitrogen. Total RNA was extracted from the frozen samples using Monarch Total RNA miniprep (New England Biolabs) and quantified using Qubit assay. Approximately 5 µg of total RNA were visualized using a 1% agarose gel supplemented with 1% bleach. The dsRNA of *C. elegans gfp* fragment was expected to be approximately 600-700 bp, and was visible without depleting bacterial ssRNAs.

#### dsRNA quantification assay

*E. coli* strains were grown overnight as described above, subcultured (1:100 dilution), and grew to an OD600 of 1.0. Cultures were divided into 1 mL aliquots, pelleted by centrifugation, flash-frozen in liquid nitrogen, and stored at −80°C until use. Total RNA was extracted and quantified as described above. Double-stranded RNA was quantified using a J2 antibody-based dsRNA ELISA kit (GenScript) according to the manufacturer’s instructions. For each sample, 10 μL of total RNA was analyzed. Briefly, a seven-step, threefold serial dilution was prepared for each RNA sample, resulting in eight assay wells, including the undiluted sample. Two biological replicates were analyzed for each strain. A standard curve was generated in GraphPad Prism using simple linear regression, and the resulting equation was used to calculate the dsRNA concentration of each sample.

To estimate dsRNA production per plasmid copy, three consecutive dilutions showing the expected approximately threefold change in measured concentration were selected to ensure measurements were within the linear range of the assay and free from saturation. The dsRNA concentration measured for each selected dilution was multiplied by its corresponding dilution factor to determine the concentration in the original sample. This procedure yielded six independent concentration estimates for each strain (three dilution points from each of two biological replicates). For background correction, the mean dsRNA concentration measured from the wild-type (no plasmid) control was calculated from the six corresponding measurements and subtracted from each individual measurement of all experimental groups. The background-corrected dsRNA concentration was then divided by the estimated plasmid copy number to determine the amount of dsRNA produced per plasmid copy.

Plasmid copy numbers were estimated from published values by assuming that an *E. coli* culture at OD600 = 1.0 contains approximately 8 × 10^8^ cells/mL. ColE1 (pUC)-based plasmids were assumed to have an average copy number of approximately 650 copies per cell, whereas L4440 plasmids were assumed to have an average copy number of approximately 75 copies per cell, based on published reports (19, 20). Background-corrected dsRNA production per plasmid copy was plotted and analyzed in GraphPad Prism using one-way analysis of variance (ANOVA).

### RNA interference in *C. elegans* by feeding bacteria

RNA interference in *C. elegans* was adapted from a published protocol. Briefly, feeding strains of *E. coli* were grown in LB liquid media supplemented with kanamycin (50 µg/mL), incubated overnight at 37℃ with aeration. Bacterial overnight cultures were concentrated by centrifugation and seeded onto NGM agar supplemented with kanamycin (50 µg/mL) and incubated at room temperature overnight. Five adult nematodes (PD4251 strain) were placed on RNAi plates for three hours to allow egg laying, then removed (Day 0), yielding ∼50–80 synchronized progeny per plate. For induction experiments, IPTG was supplemented to RNAi NGM agar plates at final concentrations of 0 mM, 1 mM, 2 mM, and 3 mM to establish a gradient of dsRNA expression. Nematode eggs were cultured on the same plates, incubated at 25℃ until confocal imaging and quantitative analysis of RNAi. To maintain IPTG stability, plates were wrapped in foil and kept in the dark. Nematodes were imaged on Days 3–5, and the optimal time point was selected based on GFP fluorescence.

### Confocal imaging and quantitative analysis

RNAi-mediated knockdown of *gfp* expression in *C. elegans* PD4251 and N2 *gfp* autofluorescence was assessed using confocal microscopy. Young adult hermaphrodite nematodes were imaged using Zeiss LSM 900. For each individual nematode, z-stack images were acquired in the GFP channel, covering the full body length and width of the animal under identical imaging conditions were held constant across all samples. Z-stack images were projected into a two-dimensional maximal intensity plot using Zeiss ZenPro Software.

Quantitative analysis of GFP fluorescence was performed using Fiji (ImageJ) based on the reconstructed z-stack projections. Briefly, each nematode was manually outlined to define a single region of interest (ROI), and fluorescence was quantified as whole-animal total GFP intensity. Integrated density was measured for each ROI, and background fluorescence was determined from an adjacent region lacking detectable GFP signal. Corrected total fluorescence (CTF) was calculated using the following equation:

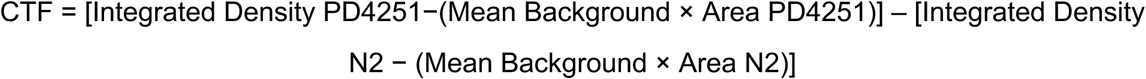

This approach enabled quantitative comparison of total GFP fluorescence at the level of individual nematodes across experimental conditions.

Statistical analysis was performed using GraphPad Prism (version 10.6.1). Differences among multiple experimental conditions were assessed using one-way analysis of variance (ANOVA), with comparisons made relative to the empty-vector control. Statistical significance was defined as *p* < 0.05.

## Acknowledgements

We thank Andrew Fire for discussing dsRNA extraction. We thank Grischa Chen for training us on quantitative confocal imaging. Margaret McFall-Ngai shared the confocal microscope. Richard Murray and Miki Yun shared BioTek plate readers and provided feedback with expertise in the CIDAR MoClo system. This work is supported by Carnegie Science endowment and Colorado College internal fund.

## Authors contributions

**MX**: promoter screen, construction of dsRNA part library, RNAi feeding assays, confocal imaging and data analysis, writing of first draft.

**SWI**: dsRNA extraction, visualization, and quantification by ELISA. Setting up RNAi feeding assay, writing first draft.

**MP**: Initiating the project, conceiving engineering ideas, establishing engineering pipeline, construction of dsRNA part library.

**PL**: Mentoring MX, writing first draft and editing, providing funding.

**MC**: Initiating and overseeing the project, conceiving engineering ideas, establishing engineering pipeline, designing and performing experiments, outlining manuscript, editing, mentoring MX, SWI, and MP, providing funding.

## Supplemental Materials

**Figure S1:**
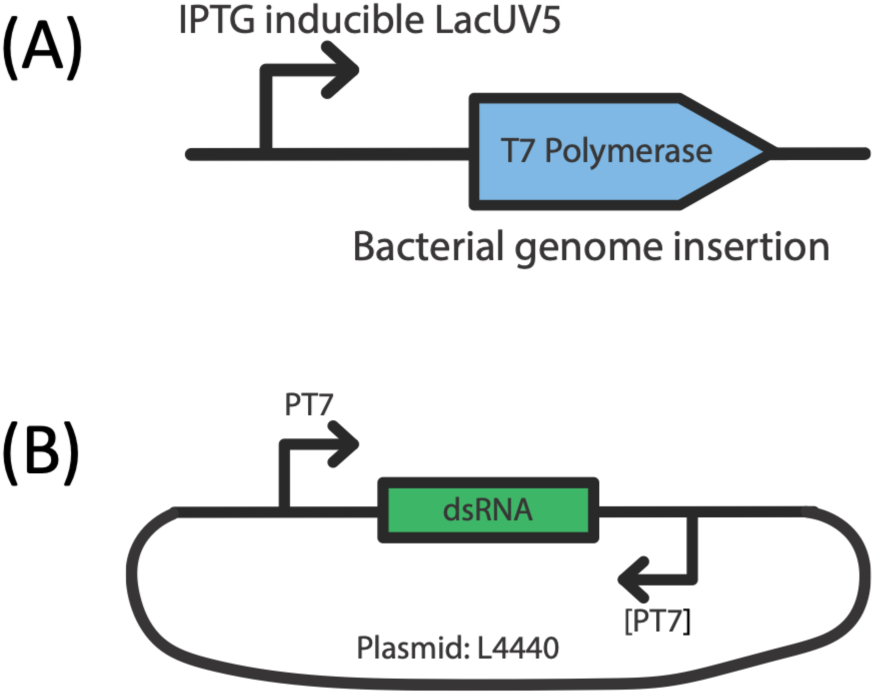
Current biocircuit architecture used for dsRNA expression in *C. elegans* RNAi includes two components. (A): a genome-integrated and IPTG-inducible promoter LacUV5 driving T7 RNA polymerase. LacUV5 is also known to be “leaky” causing T7 RNA polymearse production and downstream dsRNA expression without induction. (B): A plasmid L4440 carrying T7 promoters driving dsRNA production in the presence of T7 RNA polymerase.

**Figure S2:**
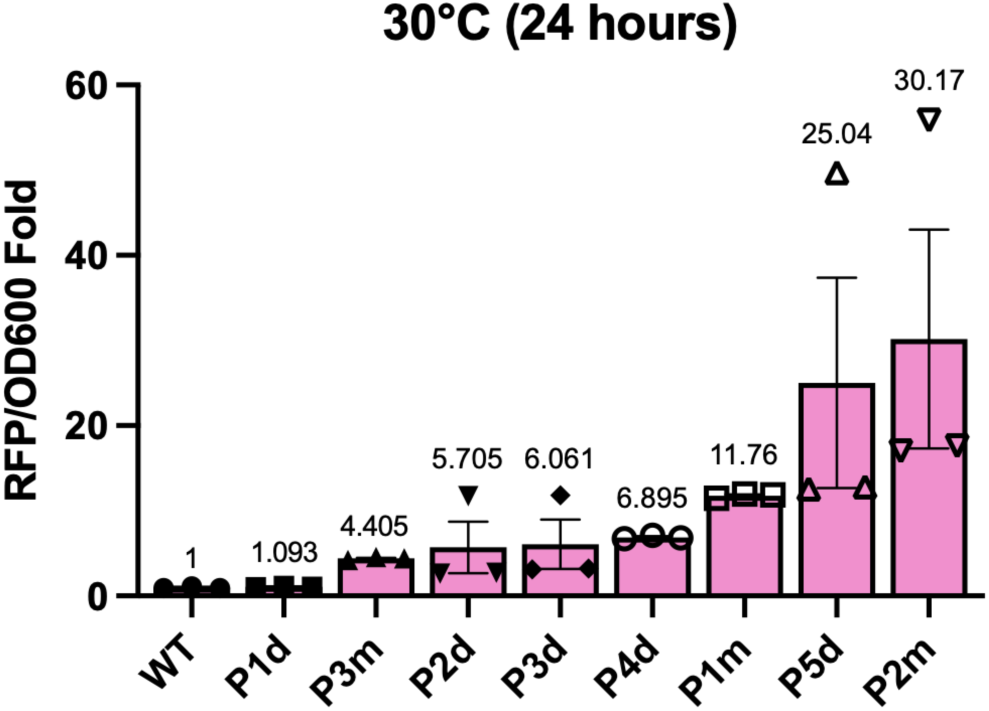
Constitutive promoter screen at 30°C in vitro. Promoter activity was assessed using RFP fluorescence intensity divided by cell density (OD600) at 24 hours growth, normalized to wild-type (WT) carrying empty vector. Each data point represents one biological replicate. Average and standard errors were shown.

**Figure S3:**
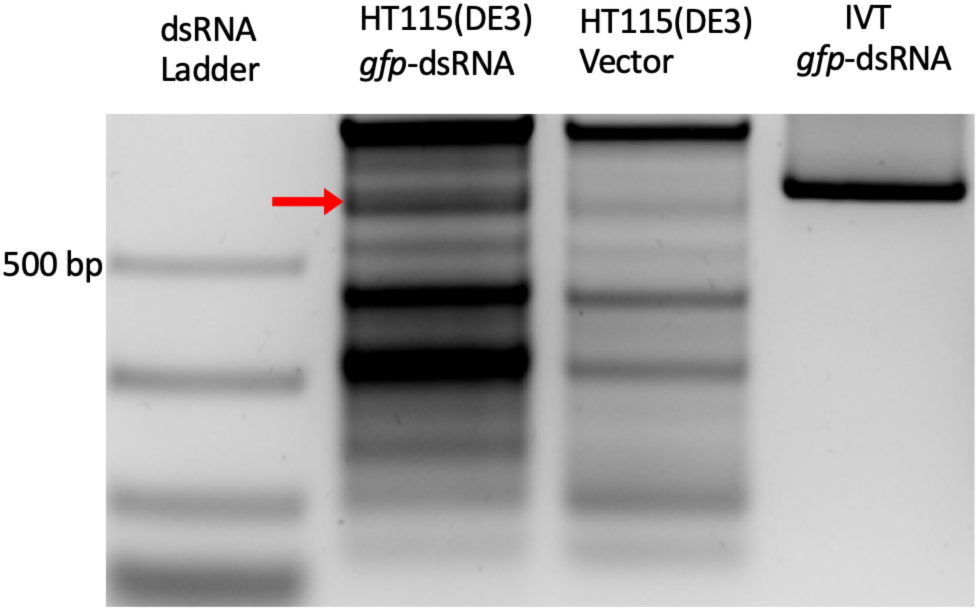
Visualization of dsRNA: 1% bleached agarose gel electrophoresis of total RNA extracted from HT115(DE3)-*gfp* strain, HT115(DE3) empty vector and *in vitro* transcribed *gfp*-dsRNA (5 µg RNA per sample). A dsRNA molecular marker was used for size estimation, and the *in vitro* transcription (IVT) *gfp-*dsRNA (739 bp), which is slightly larger than the HT115(DE3) *gfp-*dsRNA (693 bp), was included as a control. The bacterial dsRNA is indicated by a red arrow at approximately 693 bp.

**Table S1.** New part plasmids are constructed in this work*.

| Plasmid | Category | Construct | Overhangs | Description |
| --- | --- | --- | --- | --- |
| pMX001 | CDS | Partial GFP | CD | Partial L4440 C-GFP-D (643bp), BsaI cutting site removed |
| pMX002 | Reverse Promoter | Reverse Ptac | DE | Reverse complement of Ptac with DE overhangs |
| pMP001 | Reverse Terminator | Reverse T2m | AB | Reverse complement of T2m with AB overhangs |
| pMP002 | Promotor | P1m | BC | Forward B-P1m-C |
| pMP003 | Promotor | Reverse P1m | DE | Reverse complement of P1m with DE overhangs |
| pMP004 | Promotor | Ptac | BC | Forward B-Ptac-C |
| pMP005 | Terminator | T12m | EF | Forward E-T12m-F |
| pMX019 | Promotor | P2d | BC | Forward B-P2d-C |
| pMX020 | Promotor | Reverse P2d | DE | Reverse complement of P2d with DE overhangs |
| pMX021 | Promotor | P3m | BC | Forward B-P3m-C |
| pMX022 | Promotor | Reverse P3m | DE | Reverse complement of P3m with DE overhangs |
| pMX023 | Promotor | P3d | BC | Forward B-P3d-C |
| pMX024 | Promotor | Reverse P3d | DE | Reverse complement of P3d with DE overhangs |
| pMX025 | Promotor | P4d | BC | Forward B-P4d-C |
| pMX026 | Promotor | Reverse P4d | DE | Reverse complement of P4d with DE overhangs |
| pMX027 | Promotor | P5d | BC | Forward B-P5d-C |
| pMX028 | Promotor | Reverse P5d | DE | Reverse complement of P5d with DE overhangs |

\*All part plasmids are constructed based on the DVA backbone, a high-copy plasmid with Carbenicillin resistance as a selective marker.

**Table S2.** Circuit constructed for RNAi feeding and promoter screening*.

| Plasmid | Category | Construct | Description |
| --- | --- | --- | --- |
| pMX003 | Empty vector | N/A | Empty vector plasmid |
| pMX004 | RNAi feeding | UNS1-[T2m]-P1m-GFP-[P1m]-T12m-UNSX | tuMX001(GFP with J23101) |
| pMX005 | RNAi feeding | UNS1-P4d-U1d-LacI-T2m-UNS3-[T2m]-P61m-GFP-[P61m]-T12m-UNSX | IPTG inducible, LacI+GFP with Ptac |
| pMX006 | Promoter screening | UNS1-P1d-U3d, C5m-T12m-UNSX | P1d promoter screening with rfp |
| pMX008 | Promoter screening | UNS1-P2d-U3d, C5m-T12m-UNSX | P2d promoter screening with rfp |
| pMX009 | Promoter screening | UNS1-P3d-U3d, C5m-T12m-UNSX | P3d promoter screening with rfp |
| pMX010 | Promoter screening | UNS1-P3m-U3d, C5m-T12m-UNSX | P3m promoter screening with rfp |
| pMX011 | Promoter screening | UNS1-P4m-U3d, C5m-T12m-UNSX | P4m promoter screening with rfp |
| pMX012 | Promoter screening | UNS1-P4d-U3d, C5m-T12m-UNSX | P4d promoter screening with rfp |
| pMX013 | Promoter screening | UNS1-P2m-U3d, C5m-T12m-UNSX | P2m promoter screening with rfp |
| pMX014 | Promoter screening | UNS1-P1m-U3d, C5m-T12m-UNSX | P1m promoter screening with rfp |
| pMX015 | Promoter screening | UNS1-P5d-U3d, C5m-T12m-UNSX | P5d promoter screening with rfp |

\*All circuits were constructed based on the ColE1H v19d backbone, a high-copy plasmid with Kanamycin resistance as a selective marker. '[ ]' denotes reverse complement sequences.

## Notes

### Competing Interest Statement

The authors have declared no competing interest.

### Summary of Updates

In this revision, we have expanded our part library with new plasmid constructs and incorporated additional experimental validation to quantify and characterize dsRNA production with appropriate controls. We have also revised multiple sections of the manuscript to better benchmark our system against existing RNAi platforms, and to provide additional methodological details.

